# 1 Degree heating weeks fail to reach alert thresholds yet coral bleaching is widespread: structural insensitivity of anomaly-based metrics across Japan’s latitudinal gradient

**DOI:** 10.64898/2026.03.13.711501

**Authors:** Hiroki Fukui

## Abstract

Degree Heating Weeks (DHW) is the standard metric for global coral bleaching prediction, yet its performance varies markedly across regions for structurally unexplained reasons. We analyse five years (2020–2024) of standardised bleaching surveys from Japan’s Monitoring Site 1000 program (26 sites, 113 site-years; balanced panel n = 105) across a 24–35°N latitudinal gradient to diagnose why DHW fails in subtropical waters. Only 4 of 113 site-years (3.5%) reached the DHW ≥ 4 alert threshold, while bleaching (>0%) was recorded in 65 site-years (57.5%). DHW sensitivity for detecting any bleaching was 6.2%. A simple absolute-temperature metric (days with SST ≥ 30°C) significantly outperformed DHW in discriminating ≥50% bleaching (AUC = 0.926 vs 0.667, ΔAUC = 0.260, 95% CI [0.154, 0.355], p < 0.001), with the largest gap at low latitudes (ΔAUC = 0.293, p < 0.001). The Maximum Monthly Mean (MMM) was strongly correlated with latitude (r = −0.914), compressing the thermal gap available for HotSpot accumulation at low-latitude sites and eliminating HotSpot events at high-latitude sites. This structural insensitivity — arising from the anomaly-based design of DHW rather than from threshold miscalibration — operated through two distinct mechanisms across the latitudinal gradient. At low latitudes, where MMM approaches 30°C, HotSpot signals were compressed below detection thresholds despite widespread bleaching; at high latitudes, SST rarely exceeded MMM, rendering HotSpot events absent altogether. These findings demonstrate that DHW’s standard alert framework is structurally non-functional across Japan’s coral monitoring network and that regional assessment requires metrics independent of the MMM-relative anomaly architecture.

## 1.2 1. Introduction

Degree Heating Weeks (DHW) underpins the global coral bleaching monitoring infrastructure operated by NOAA Coral Reef Watch. Computed as the cumulative positive anomaly of sea surface temperature (SST) above a site’s Maximum Monthly Mean (MMM) over a trailing accumulation window — conventionally 12 weeks in NOAA’s operational product (Liu et al. 2014), though Lachs et al. (2021) demonstrated that an 8-week window optimises bleaching prediction — DHW translates thermal stress into a single metric with standardised alert thresholds: DHW ≥ 4 °C-weeks signals significant bleaching likely, and DHW ≥ 8 °C-weeks signals mass bleaching and mortality likely (Liu et al. 2014; Skirving et al. 2020). Since its operational deployment, DHW has served as the primary basis for bleaching early warnings, reef management decisions, and retrospective attribution of bleaching events to thermal stress (Eakin et al. 2010), including the recently declared fourth global mass bleaching event (Spady et al. 2026). The central design principle of DHW is its anomaly-based architecture: rather than measuring absolute temperature, it measures how far and for how long SST departs from the local climatological maximum. This relativistic design was intended to make DHW universally applicable — a single metric that functions identically whether applied to equatorial, subtropical, or marginal reef environments.

At regional scales, however, DHW’s predictive performance is far from uniform. Whitaker and DeCarlo (2024) demonstrated that regionally optimising the DHW definition — adjusting parameters such as the HotSpot threshold and accumulation window — substantially improved bleaching prediction skill across diverse reef regions. Their finding that no single set of DHW parameters performs optimally everywhere raises a question that their study did not address: what structural property of DHW necessitates this regional tuning? Lachs et al. (2021) showed that the choice of MMM baseline period and the HotSpot threshold filter (MMM+0°C vs MMM+1°C) significantly affect predictive accuracy, further demonstrating that DHW performance is sensitive to internal parameter choices. Together, these studies establish that DHW requires regional calibration but do not explain why. A temporal dimension of this problem has been explored by Lachs et al. (2023), who simulated coral thermal tolerance increases at Palau and showed that progressive upward shifts in the MMM baseline erode the DHW signal over decadal timescales. However, the spatial mechanism — how the absolute value of MMM constrains DHW signal range across contemporary reef environments — has not been identified.

A recent analysis of Japan’s Monitoring Site 1000 data provided an empirical clue. Fukui (2026) analysed five years of standardised bleaching surveys across 26 sites spanning 24–35°N and found that a simple absolute-temperature metric — the number of days with SST ≥ 30°C — substantially outperformed DHW in discriminating bleaching across all severity levels (AUC = 0.877 vs 0.624 at ≥50% prevalence, p < 0.001). Where that study documented *what* predicts bleaching and characterised the distributional structure of bleaching prevalence, the present study addresses the logically prior question of *why* the standard monitoring metric fails — a question whose answer has direct consequences for whether and how DHW can be reformed for regional application. The present study asks: why does DHW fail to predict coral bleaching in Japan’s subtropical monitoring network, and does the failure mechanism differ across latitudes?

Japan’s Monitoring Site 1000 program provides an unusually well-suited system for addressing this question. The network spans approximately 10° of latitude (24–35°N) under a single standardised protocol, from the Yaeyama Islands — where MMM approaches 30°C and reefs experience chronic high temperatures — to Tateyama Bay near the northern limit of hermatypic corals in the northwestern Pacific, where MMM is approximately 27°C. This latitudinal gradient creates a natural experiment in which the MMM — the baseline upon which DHW’s entire anomaly architecture rests — varies systematically. If DHW’s anomaly-based design interacts with the absolute value of MMM to produce regionally variable performance, this gradient should reveal both the interaction and its mechanism. We approach this question through three complementary analyses: first, we evaluate the diagnostic performance of standard DHW alert thresholds (≥4, ≥8 °C-weeks) across latitudinal bands; second, we compare the continuous discriminatory performance (AUC) of DHW and an absolute-temperature metric across latitudes; and third, we characterise the thermal safety margin — the gap between MMM and observed summer maximum SST — to identify the structural mechanism underlying DHW’s variable performance.

## 1.3 2. Methods

### 1.3.1 2.1 Study system and data

Coral bleaching data were obtained from the Monitoring Site 1000 coral reef survey program, administered by the Ministry of the Environment, Japan (data citation: Ministry of the Environment, Japan. Monitoring Site 1000 Coral Reef Survey, fiscal years 2020–2024; approved under permit Kansei-Tahatsu No. 2602271). The program employs a standardised spot-check protocol across sites spanning the Ryukyu Archipelago, Ogasawara Islands, and adjacent temperate waters. The dataset comprises 585 survey points at 26 sites, covering a latitudinal range of 24.26–34.98°N — from the Yaeyama Islands in the southern Ryukyus to Tateyama Bay near the northern limit of hermatypic corals in the northwestern Pacific — and yielding 113 site-years of observations over five fiscal years (2020–2024). The five-year period encompasses two major bleaching events (2022 and 2024) and three milder years (2020, 2021, 2023).

Bleaching prevalence was recorded at each survey point using a 1 m × 1 m line-intercept belt transect method, in which the ratio of bleached colony area to total colony area was quantified. Surveys are conducted annually in autumn (September–November), approximately 1–3 months after the summer thermal stress peak (see Fukui 2026 for full protocol details). For the present analysis, point-level observations were aggregated to the site level by taking the median bleaching prevalence across all survey points within each site × fiscal year combination (n = 113 site-years across 26 sites). A balanced panel of 21 sites observed in all five fiscal years (n = 105 site-years) was used for all comparative and inferential analyses to avoid compositional bias; five sites (02, 08, 20, 25, 26) had fewer than five years of data and were included only in descriptive summaries. The full dataset (n = 113) was used for threshold classification (Section 2.5) and distributional summaries.

### 1.3.2 2.2 Satellite sea surface temperature

Daily sea surface temperature (SST) was obtained from the Multi-scale Ultra-high Resolution SST (MUR SST v4.1; JPL MUR MEaSUREs Project 2015), which provides global coverage at 0.01° (∼1 km) resolution. SST values were extracted for each survey point using the nearest MUR grid cell. For the 585 survey points, SST was available for 2019–2024; gaps at individual points (due to land masking or cloud contamination) were filled using the site-level daily mean SST. Within-site SST standard deviations ranged from 0.17 to 0.48°C.

### 1.3.3 2.3 Thermal stress metrics

Two thermal stress metrics were computed from MUR SST, so that differences in predictive performance reflect metric structure rather than input data.

#### Degree Heating Weeks (DHW; Lachs method)

The Maximum Monthly Mean (MMM) was calculated as the maximum of monthly mean SSTs over the 2019–2024 period for each site. HotSpots were defined as max(SST − MMM, 0). DHW was computed as the cumulative HotSpot over a trailing 56-day (8-week) window, divided by 7, yielding units of °C-weeks. This procedure follows Lachs et al. (2021), who demonstrated that accumulating all positive anomalies above MMM (without an additional threshold filter) improves bleaching prediction relative to the standard NOAA approach. Our implementation differs from that of Lachs et al. (2021) in the climatological baseline: we used 2019–2024 MUR SST rather than the 1985–1993 fixed climatology, because the short and recent observational period precluded construction of a multi-decadal baseline independent of the study interval. This choice means that the MMM reflects contemporary thermal regimes, including any warming trend over 2019–2024; the implications are addressed in the Discussion.

#### DHW (NOAA method; sensitivity analysis)

For comparison with the standard operational product, we also computed DHW using the NOAA Coral Reef Watch definition, in which HotSpots are defined as max(SST − (MMM + 1°C), 0) (Liu et al. 2014). All other parameters (56-day accumulation window, division by 7) were identical to the Lachs method.

#### Threshold exceedance days (days30)

Defined as the number of days with SST ≥ 30°C during the 90-day period preceding each site’s survey date. This metric was identified as the optimal absolute-threshold predictor of bleaching in Fukui (2026).

### 1.3.4 2.4 Latitudinal classification

Sites were assigned to three latitudinal bands following Fukui (2026): low (≤26°N; predominantly the Yaeyama Islands), mid (26–30°N; Okinawa main island to Ogasawara), and high (>30°N; Yakushima to Tateyama). In the balanced panel, the low band contained 9 sites (45 site-years), the mid band 6 sites (30 site-years), and the high band 6 sites (30 site-years).

### 1.3.5 2.5 DHW threshold analysis

The conventional DHW alert framework classifies thermal stress by fixed thresholds: DHW ≥ 4 indicates significant bleaching likely, and DHW ≥ 8 indicates widespread bleaching and mortality likely (Liu et al. 2014). To evaluate the diagnostic utility of these thresholds across latitudinal bands, we classified each site-year as a true positive, false positive, false negative, or true negative at DHW thresholds of 3, 4, 5, 6, 7, and 8 °C-weeks. Three binary bleaching response definitions were used: any bleaching (>0%), moderate-to-severe bleaching (≥50%), and severe bleaching (≥80%). False positive rate (FPR), false negative rate (FNR), accuracy, sensitivity, and specificity were computed for each threshold × response × latitudinal band combination.

### 1.3.6 2.6 Predictive performance comparison

Discriminatory performance of DHW and days30 was compared using the area under the receiver operating characteristic curve (AUC). To account for within-site temporal correlation, AUC values were derived from generalised estimating equations (GEE; binomial family, logit link, exchangeable correlation structure) with site as the clustering unit. The difference in AUC between metrics (ΔAUC = AUC_days30 − AUC_DHW) was tested using a cluster bootstrap procedure: sites (the clustering unit) were resampled with replacement (1,000 iterations), and ΔAUC was computed for each bootstrap sample. The 95% confidence interval was obtained by the percentile method, and the p-value was computed as the proportion of bootstrap samples in which ΔAUC ≤ 0. This site-level resampling preserves the within-site temporal correlation that conventional methods such as DeLong’s test (DeLong et al. 1988) do not accommodate.

To test whether the DHW–bleaching association varied with latitude, we fitted GEE interaction models of the form bleach ∼ predictor × latitude_band, where bleach was the binary bleaching outcome (≥50%) and predictor was either DHW or days30. Non-significance of the interaction term would indicate that the latitudinal gradient in AUC is not attributable to a differential slope of the predictor–bleaching relationship.

### 1.3.7 2.7 Thermal safety margin analysis

For each site, we computed the MMM and the median summer maximum SST (daily maximum during July–September over 2020–2024). The thermal gap was defined as the difference between the summer maximum SST and MMM. This quantity represents the maximum anomaly a site can generate under observed conditions and therefore sets an upper bound on HotSpot accumulation. MMM was regressed against latitude (Pearson’s r). The thermal gap analysis is descriptive; to avoid circularity, thermal gap was not included as a predictor in the AUC models.

### 1.3.8 2.8 Sensitivity analyses

Three sensitivity analyses were performed. First, AUC values were recomputed using the NOAA-standard DHW (with MMM + 1°C baseline filter) alongside the Lachs-method DHW and days30. Second, a leave-one-year-out procedure was applied: each of the five fiscal years was excluded in turn, and ΔAUC was recomputed on the remaining four years. This procedure tests whether the overall advantage of days30 over DHW is driven by a single anomalous year. Third, to assess whether the MUR-derived DHW values used in this study are consistent with the operational NOAA CRW product, we compared annual maximum DHW values at each site for 2020–2024 between our MUR-based calculation (Lachs method, 8-week window, MMM derived from 2019–2024) and the NOAA CRW Daily Global 5km product (v3.1; 12-week window, MMM+1°C threshold, 1985–2012 climatology; Liu et al. 2014). CRW DHW values were extracted from the NOAA CRW ERDDAP server (NOAA Coral Reef Watch 2026) for the nearest 5km pixel to each site, covering July–October of each year.

All analyses were conducted in Python 3.11 using pandas, scipy, numpy, statsmodels, and matplotlib.

## 1.4 3. Results

### 1.4.1 3.1 DHW rarely reaches alert thresholds

Across all 113 site-years, median DHW was 0.041 °C-weeks (IQR: 0.000–0.507), and only 4 site-years (3.5%) reached the DHW ≥ 4 alert threshold (Fig. 1, Table 1). Of these four, three occurred in the mid-latitude band and one in the low-latitude band; no high-latitude site-year reached DHW ≥ 4. DHW distribution statistics varied markedly across bands: in the low-latitude band, median DHW was 0.064 °C-weeks (IQR: 0.000–0.332), with only 2.1% of site-years reaching DHW ≥ 4 despite 85.1% recording at least one day above 30°C. In the mid-latitude band, median DHW was higher (0.495 °C-weeks; IQR: 0.000–1.835) and 9.7% of site-years exceeded DHW ≥ 4. In the high-latitude band, the maximum observed DHW was 2.56 °C-weeks and no site-year exceeded even DHW ≥ 3; median DHW was 0.000.

**Fig. 1.**
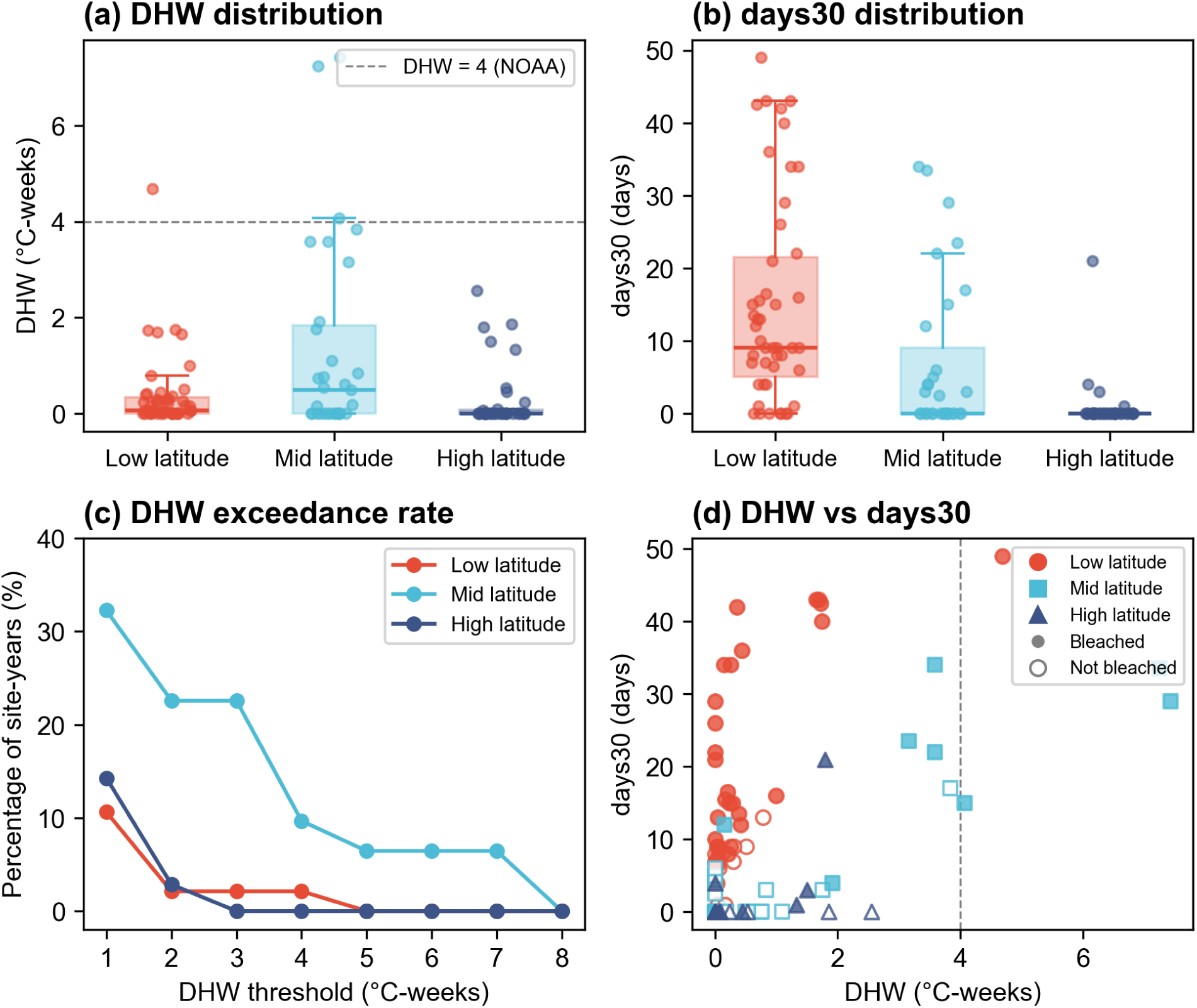
Distribution of DHW values across 113 site-years, contrasted with bleaching occurrence. Dashed line indicates DHW = 4 alert threshold.

**Table 1.**
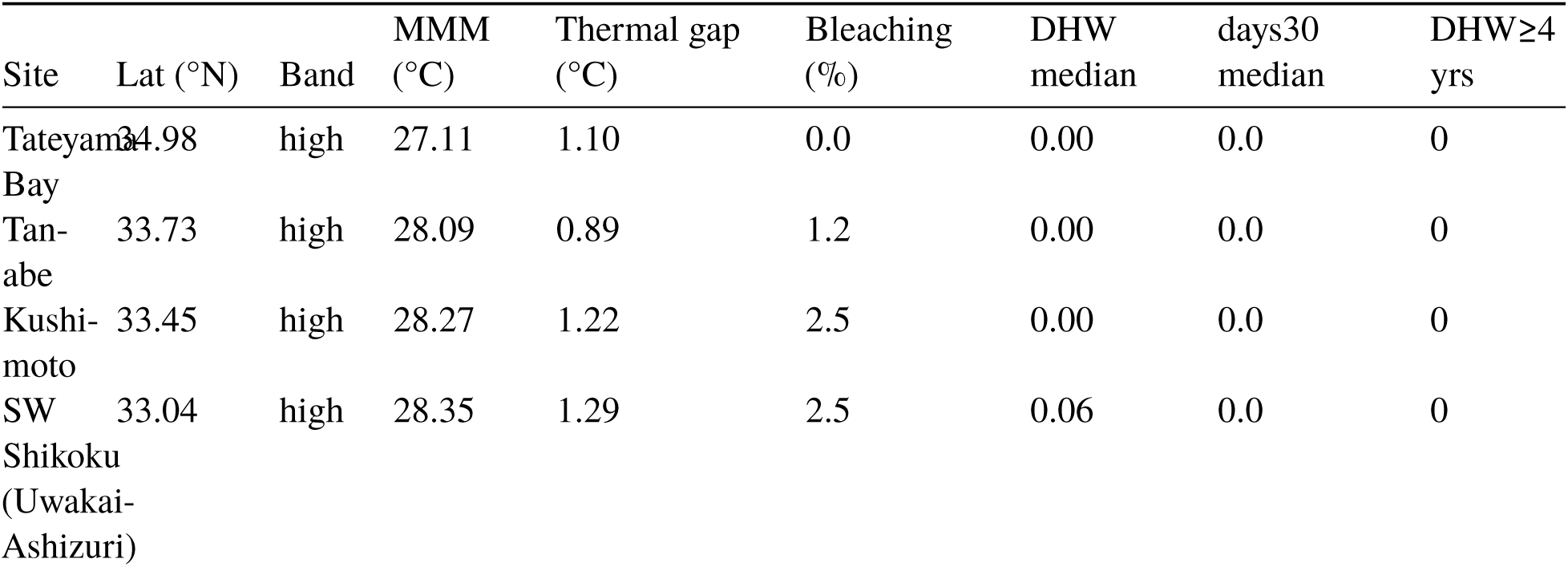
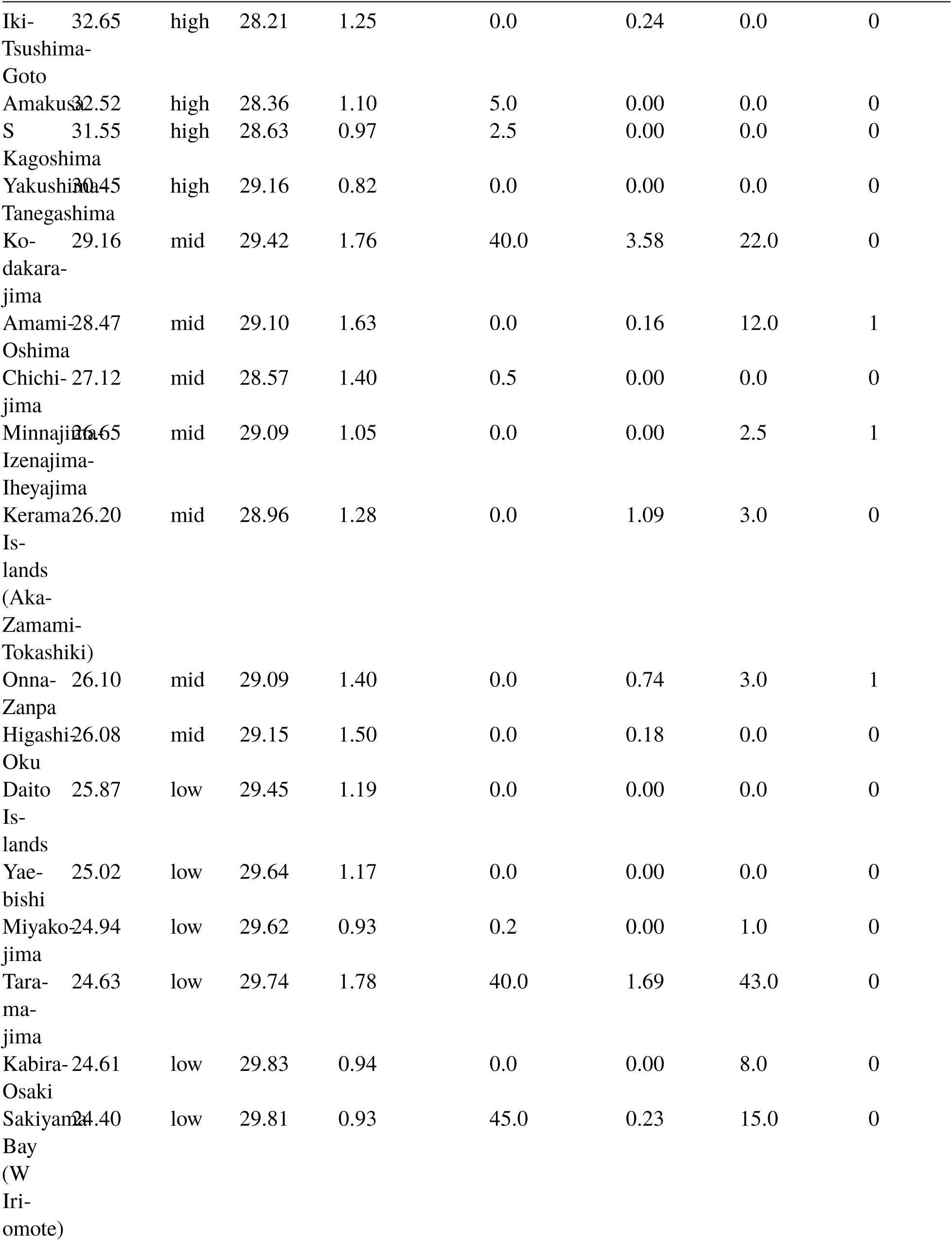
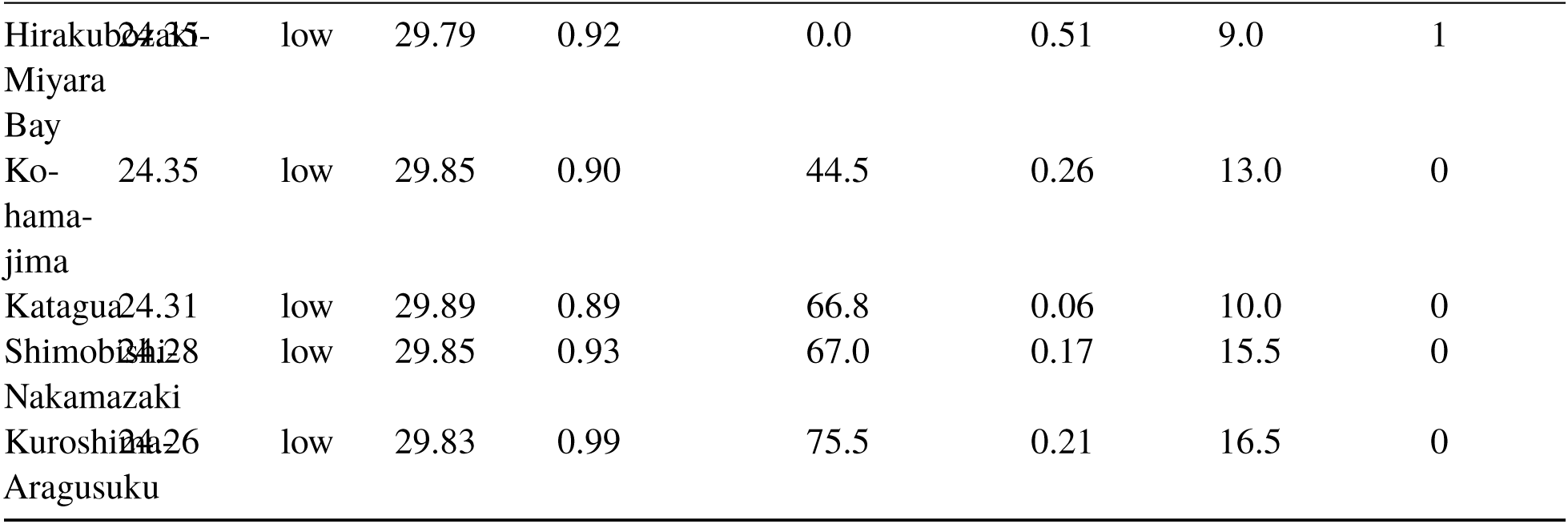
Site-level summary of thermal metrics and bleaching across the latitudinal gradient.

In contrast, bleaching (>0%) was recorded in 65 of 113 site-years (57.5%), and ≥50% bleaching occurred in 26 of 113 site-years (23.0%). At the conventional DHW ≥ 4 alert threshold applied to any bleaching (>0%), sensitivity was 6.2% (4/65), with a false negative rate of 93.8%. For ≥50% bleaching, DHW ≥ 4 detected only 2 of 26 events (sensitivity = 7.7%) while generating 2 false positives (Table 1). The days30 metric showed substantially greater signal range: the median was 1.0 day (IQR: 0.0–13.0), with 52.2% of site-years recording at least one day above 30°C and 46.0% recording three or more days.

### 1.4.2 3.2 Predictive performance degrades at low latitudes

Using the ≥50% bleaching threshold as the primary outcome (prevalence = 24.8% in the balanced panel), days30 significantly outperformed DHW overall (AUC = 0.926 vs 0.667; ΔAUC = 0.260, 95% CI [0.154, 0.355], p < 0.001; Table 2, Fig. 3).

**Fig. 2.**
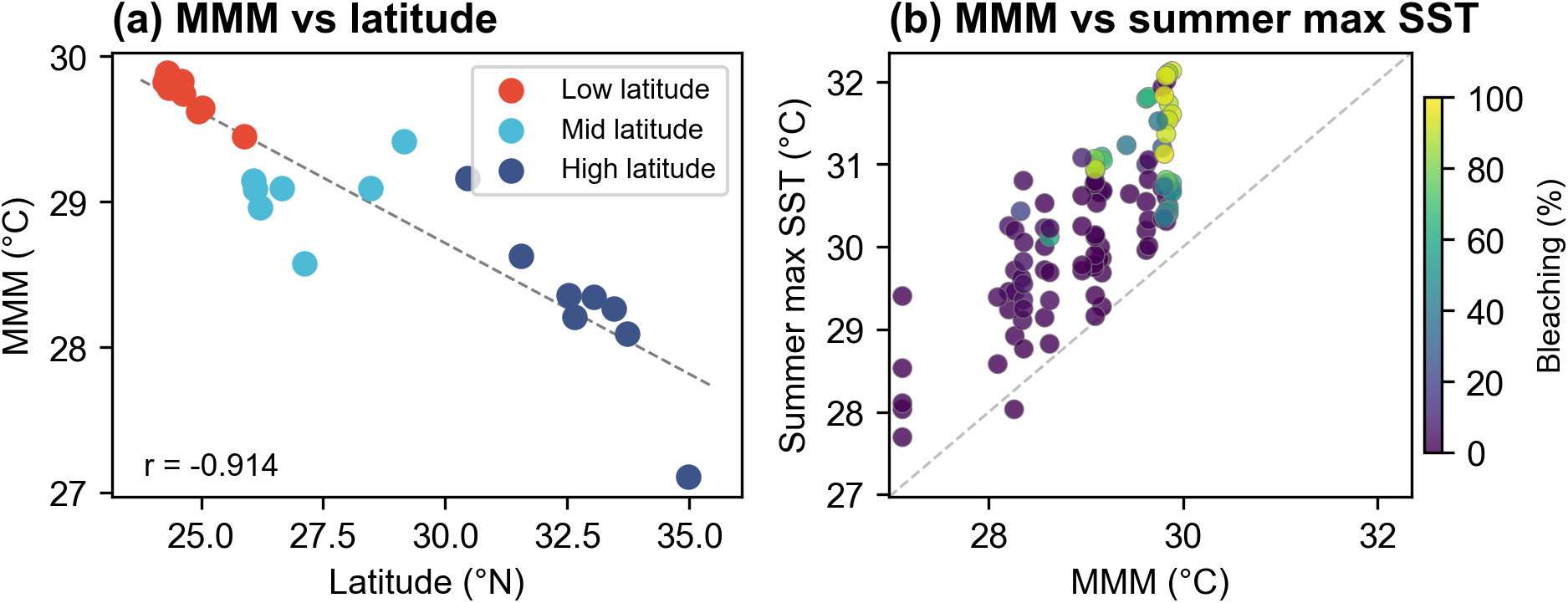
Relationship between MMM and latitude across 26 sites, with thermal gap (summer max SST − MMM) by latitudinal band. (a) MMM versus latitude with linear regression. (b) Summer maximum SST versus MMM for each site-year, coloured by bleaching prevalence. Dashed diagonal line indicates the 1:1 relationship (thermal gap = 0), where summer maximum SST equals MMM.

**Fig. 3.**
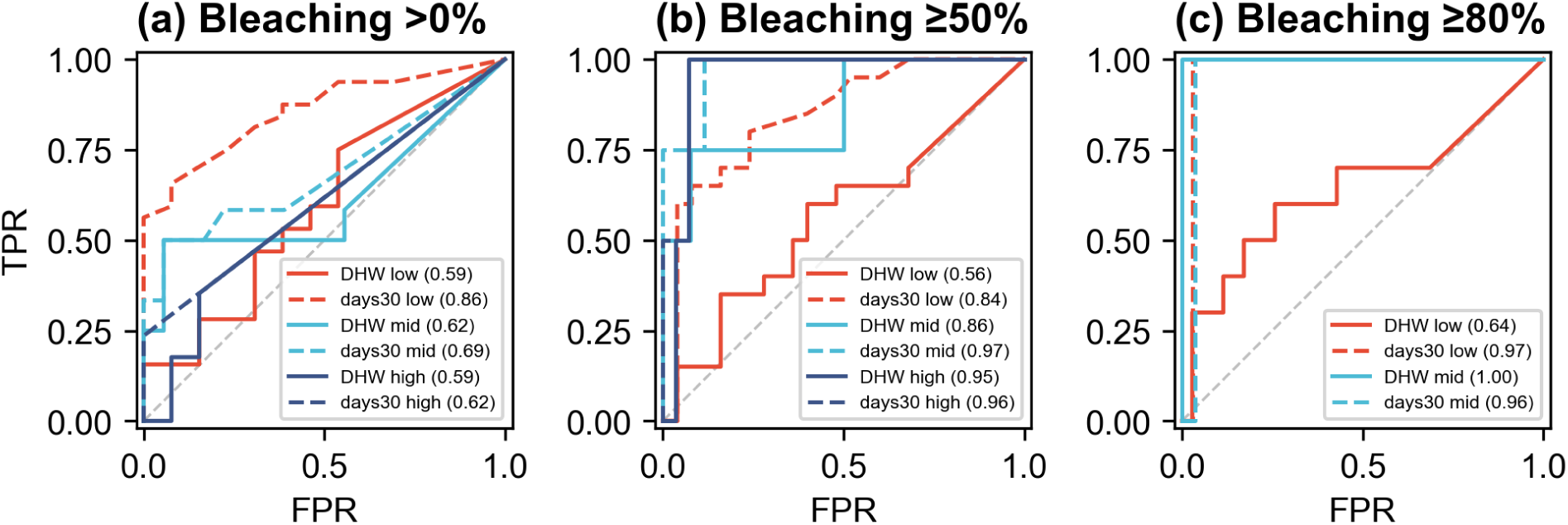
ROC curves for DHW and days30 by latitudinal band at the ≥50% bleaching threshold.

**Table 2.** AUC comparison between DHW and days30 by bleaching threshold and latitudinal band (balanced panel, n = 105).

| Bleach thresh | Lat band | n | DHW AUC | days30 AUC | ΔAUC | 95% CI | p |
| --- | --- | --- | --- | --- | --- | --- | --- |
| >0% | all | 105 | 0.571 | 0.735 | 0.161 | [0.036, 0.274] | 0.003 |
| >0% | low | 45 | 0.594 | 0.856 | 0.258 | [0.196, 0.364] | 0.000 |
| >0% | mid | 30 | 0.616 | 0.685 | 0.068 | [-0.022, 0.145] | 0.097 |
| >0% | high | 30 | 0.586 | 0.618 | 0.029 | [-0.100, 0.122] | 0.332 |
| $\geq 50\%$ | all | 105 | 0.667 | 0.926 | 0.260 | [0.154, 0.355] | 0.000 |
| $\geq 50\%$ | low | 45 | 0.562 | 0.845 | 0.293 | [0.188, 0.403] | 0.000 |
| $\geq 50\%$ | mid | 30 | 0.856 | 0.971 | 0.111 | [0.000, 0.250] | 0.333 |
| $\geq 50\%$ | high | 30 | 0.946 | 0.964 | 0.018 | [-0.069, 0.103] | 0.419 |
| $\geq 80\%$ | all | 105 | 0.687 | 0.983 | 0.297 | [0.129, 0.415] | 0.009 |
| $\geq 80\%$ | low | 45 | 0.641 | 0.971 | 0.340 | [0.174, 0.455] | 0.000 |
| $\geq 80\%$ | mid | 30 | 1.000 | 0.964 | -0.036 | [-0.103, 0.000] | 0.986 |
| $\geq 80\%$ | high | 30 | nan | nan | nan | [nan, nan] | nan |

The performance gap was largest in the low-latitude band, where DHW was near-uninformative: AUC_DHW = 0.562 versus AUC_days30 = 0.845 (ΔAUC = 0.293, 95% CI [0.188, 0.403], p < 0.001). In the mid-latitude band, DHW performed moderately well (AUC = 0.856), but days30 still achieved higher discrimination (AUC = 0.971; ΔAUC = 0.111, 95% CI [0.000, 0.250], p = 0.333). In the high-latitude band, both metrics performed well and the difference was negligible (AUC_DHW = 0.946, AUC_days30 = 0.964; ΔAUC = 0.018, 95% CI [−0.069, 0.103], p = 0.419).

For any bleaching (>0%), the overall ΔAUC was 0.161 (95% CI [0.036, 0.274], p = 0.003), with the largest gap again at low latitudes (ΔAUC = 0.258, 95% CI [0.196, 0.364], p < 0.001). At this lenient threshold, however, the high baseline prevalence in the low-latitude band (71.1%) limits the discriminatory ceiling for both metrics: even a random classifier would achieve moderate apparent performance. The mid-latitude band showed a marginally non-significant advantage for days30 (ΔAUC = 0.068, 95% CI [−0.022, 0.145], p = 0.097), while the high-latitude band showed no significant difference (ΔAUC = 0.029, 95% CI [−0.100, 0.122], p = 0.332). For severe bleaching (≥80%), the advantage of days30 was most pronounced overall (ΔAUC = 0.297, 95% CI [0.129, 0.415], p = 0.009), driven by the low-latitude band (ΔAUC = 0.340, 95% CI [0.174, 0.455], p < 0.001). The ≥80% threshold could not be evaluated in the high-latitude band because no site-year reached this severity level.

The GEE interaction models showed non-significant predictor × latitude interactions for both DHW (z = −1.33, p = 0.183) and days30 (z = −0.51, p = 0.607), indicating that the slope of the predictor–bleaching relationship does not differ significantly across latitudinal bands. The latitudinal gradient in AUC is consistent with differences in signal range rather than in ecological sensitivity, though the non-significant interaction should be interpreted with caution given the limited sample size (n = 105 site-years across 21 sites).

### 1.4.3 3.3 MMM constrains DHW signal range

MMM was strongly negatively correlated with latitude (Pearson’s r = −0.914, n = 26 sites; Fig. 2). At low-latitude sites, MMM values ranged from 29.45 to 29.89°C (median = 29.81°C), leaving a narrow thermal gap — the difference between the median summer maximum SST and MMM — of only 0.928°C (median across 11 sites). At mid-latitude sites, MMM was lower (median = 29.09°C) and the thermal gap was substantially wider (median = 1.405°C). At high-latitude sites, MMM ranged from 27.11 to 29.16°C (median = 28.31°C), with a thermal gap of 1.100°C — intermediate between the low and mid bands but generated from a much lower SST baseline.

The correspondence between thermal gap and DHW performance was monotonic across latitudinal bands: the low-latitude band, with the narrowest median thermal gap (0.928°C), produced the lowest DHW AUC (0.562); the mid-latitude band (thermal gap = 1.405°C) yielded moderate DHW performance (AUC = 0.856); and the high-latitude band (thermal gap = 1.100°C) showed the highest DHW AUC (0.946), though this largely reflected the low base rate of ≥50% bleaching (6.7%) rather than genuine discriminatory power. At the site level, the relationship between thermal gap and maximum observed DHW manifested primarily as an asymmetric ceiling constraint (Fig. S1): The 1.0°C boundary was selected a priori to approximate the median thermal gap in the low-latitude band (0.928°C), where DHW signal suppression was most severe. Of 11 sites with thermal gap < 1.0°C, only 1 (9.1%) ever reached DHW ≥ 2 over five years; in contrast, 7 of 15 sites (46.7%) with thermal gap ≥ 1.0°C exceeded this level (Fisher’s exact p = 0.084). This pattern confirms that the thermal gap sets a physical ceiling on HotSpot accumulation — narrow gaps preclude high DHW values, though wide gaps do not guarantee them because DHW also depends on the duration of exceedance.

The narrow thermal gap at low latitudes constrains the maximum HotSpot that can accumulate during a summer. Even during the severe 2024 event, when summer maximum SSTs at Yaeyama Islands sites exceeded 32°C, the resulting daily HotSpots (SST − MMM ≈ 2.0–2.3°C) were modest in magnitude and accumulated over too few consecutive days to push DHW above the ≥4 alert threshold at most sites. The low-latitude band recorded only one site-year with DHW ≥ 4 (site 11 in 2022, DHW = 4.68) despite 20 of 45 site-years (44.4%) exhibiting ≥50% bleaching (Fig. 4). By contrast, days30 registered a strong signal at these same sites: the low-latitude median was 9.0 days, with 80.9% of site-years recording ≥2 days above 30°C.

**Fig. 4.**
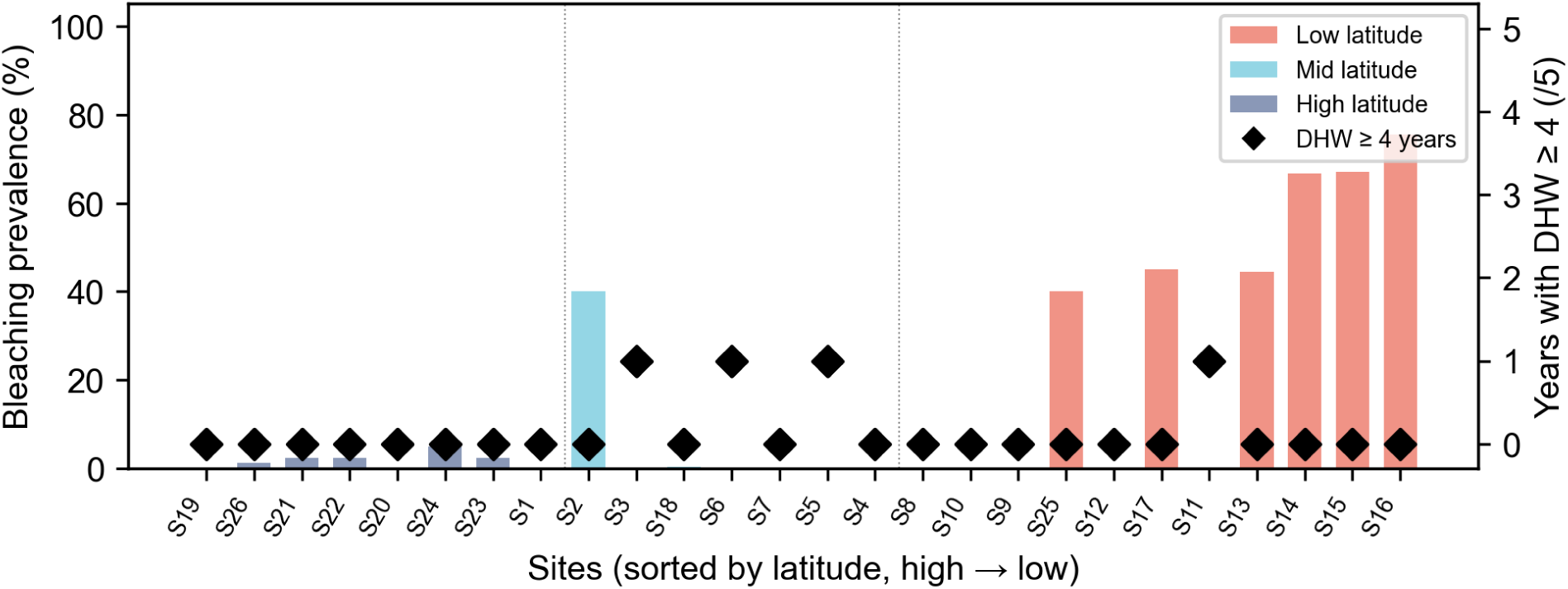
Site-level contrast between DHW and bleaching prevalence in the low-latitude band, illustrating the disconnect between the alert metric and observed bleaching. Sites are ordered by latitude (south to north) and labeled by site number (see Table 1 for site names and coordinates).

At high-latitude sites, DHW failure arose from a different mechanism: SST rarely exceeded MMM, so HotSpot events themselves were infrequent rather than merely compressed in magnitude. The maximum observed DHW at any high-latitude site was 2.56 °C-weeks, and no site-year exceeded even DHW ≥ 3 (Table 1). These two contrasting patterns — suppressed HotSpot magnitude at low latitudes and absent HotSpot events at high latitudes — suggest that DHW fails through distinct mechanisms across the latitudinal gradient.

### 1.4.4 3.4 Sensitivity analyses confirm robustness

#### NOAA-standard DHW

Replacing the Lachs-method DHW with NOAA-standard DHW (MMM + 1°C baseline filter) yielded a somewhat higher overall AUC (0.726 vs 0.667 at the ≥50% threshold) and a notable improvement in the low-latitude band (0.742 vs 0.562). This counterintuitive result — that adding a stricter filter improved discrimination — likely reflects that the +1°C offset reduces noise from small, ecologically inconsequential anomalies that dilute the DHW signal. However, days30 still substantially outperformed both DHW variants (AUC = 0.926 overall, 0.845 at low latitudes). Critically, the NOAA method’s +1°C baseline filter, applied to sites where MMM already approaches 30°C, effectively eliminates all HotSpot accumulation: no site-year reached DHW ≥ 4 under the NOAA definition, rendering the standard alert framework entirely non-functional across this monitoring network.

#### NOAA CRW operational DHW

Annual maximum DHW from the NOAA CRW 5km product was positively but weakly correlated with MUR-derived DHW (Pearson’s r = 0.351, p < 0.001; Spearman’s ρ = 0.059, p = 0.54; n = 111 site-years; Fig. S2). The divergence between Pearson and Spearman statistics reflects a large cluster of site-years in which MUR-derived DHW was near zero while CRW DHW was substantially positive, violating the monotonic relationship assumed by rank correlation. CRW DHW values were systematically higher than MUR-derived values (median MUR/CRW ratio = 0.170 among nonzero pairs), reflecting the lower MMM baseline derived from the 1985–2012 climatology and the longer 12-week accumulation window used by CRW. Critically, 56 of 111 site-years (50.5%) reached CRW DHW ≥ 4, compared with only 4 (3.6%) under the MUR-derived calculation. This stark contrast demonstrates that the structural insensitivity documented in this study — in which nearly no site-years reach the DHW ≥ 4 alert threshold — is a consequence of the contemporary (2019–2024) MMM baseline, not of the DHW framework per se. The CRW product, anchored to a pre-warming baseline, retains sufficient dynamic range to generate alerts; our MUR-based DHW, calibrated to the current thermal regime, does not. This comparison confirms the central mechanistic argument of this study: as MMM rises with climate change, the thermal gap narrows and DHW alert thresholds become progressively unreachable — a trajectory that the CRW product will itself increasingly follow as its climatology ages relative to ongoing warming.

#### Leave-one-year-out

ΔAUC (days30 minus DHW) remained positive in all five folds at the ≥50% threshold, both overall (range: 0.208–0.340) and within the low-latitude band (range: 0.154–0.386). The largest overall ΔAUC (0.340) occurred when 2024 was excluded, indicating that the advantage of days30 over DHW does not depend on the most severe bleaching year; rather, excluding 2024 amplified the contrast because DHW was particularly uninformative during the mild-to-moderate years. Within the low-latitude band, the largest ΔAUC (0.386) also occurred in the fold excluding 2024, and the smallest (0.154) when 2022 was excluded — the only year in which a low-latitude site (site 11) generated DHW ≥ 4.

## 1.5 4. Discussion

### 1.5.1 4.1 The structural insensitivity of DHW

The central finding of this study is that DHW’s failure in Japan’s monitoring network is not a problem of threshold calibration — of whether bleaching begins at DHW 4, 3, or some other value — but a consequence of the anomaly-based design itself. When MMM is high, the dynamic range of HotSpot signals is structurally compressed: SST must exceed an already-elevated baseline to generate any positive anomaly, and the magnitude of anomalies that can accumulate within a realistic summer is too small to push DHW above alert thresholds. At low-latitude sites in this study, where median MMM was 29.81°C, only one site-year reached DHW ≥ 4 despite 44.4% exhibiting ≥50% bleaching. This is not a case where DHW predicted incorrectly — it is a case where the signal that DHW is designed to measure has effectively disappeared.

Fukui (2026) noted that anomaly-based metrics measure departure from baseline whereas absolute-temperature metrics measure proximity to a physiological failure point, and that these two quantities need not be correlated. The present study extends that observation by identifying the structural mechanism: the MMM itself determines the information content of the DHW signal. Where MMM is high, DHW approaches a ceiling — not because thermal stress is absent, but because the metric’s reference frame absorbs the stress into its baseline. The metric is, in effect, calibrated against the very conditions it needs to detect.

DHW was developed by NOAA Coral Reef Watch in the 1990s and has since become the global standard for coral bleaching early warning, achieving well-documented success in detecting mass bleaching events at the pan-tropical scale (Liu et al. 2014; Skirving et al. 2020). This global-scale operational success has, however, discouraged systematic scrutiny of the metric’s regional failure modes. When a metric works well across most of its operational domain, the regions where it fails tend to be treated as local calibration problems rather than as symptoms of a structural limitation inherent in the metric’s design.

The failure pattern documented here — a baseline-relative metric losing discriminatory power as the baseline approaches the critical range — is well characterised in clinical diagnostic settings. Screening instruments calibrated against population norms routinely exhibit ceiling effects in subpopulations whose baseline scores approach the instrument’s upper bound, producing systematic false negatives in precisely the groups at highest risk. The structural parallel suggests that DHW’s regional failure is not an anomaly unique to coral reef science but rather an instance of a broader class of metrological limitations that arise whenever anomaly-based indices are applied across populations with heterogeneous baselines. That this pattern has not previously been formally diagnosed for DHW likely reflects the metric’s operational success at the global scale, which has discouraged scrutiny of its regional failure modes.

Although the arithmetic relationship between MMM and HotSpot range is in principle predictable, the present study provides the first empirical quantification of how this constraint translates into band-specific diagnostic failure — identifying both the magnitude of AUC degradation and the distinct failure mechanisms operating at low versus high latitudes — using standardised monitoring data across a continuous latitudinal gradient.

Because the thermal gap is derived directly from MMM, it cannot be included as a predictor in models alongside DHW without introducing circularity and collinearity. However, the descriptive correspondence between thermal gap and DHW performance across latitudinal bands (Section 3.3) serves a different inferential purpose: it provides a structural explanation for why the DHW signal degrades, rather than a predictive model of bleaching. This distinction — between explaining signal loss and predicting bleaching — is important for correctly interpreting the thermal gap analysis. The site-level scatter (Fig. S1) provides direct evidence that thermal gap constrains the physical ceiling of DHW, independently of bleaching outcomes. Because this relationship reflects a mathematical property of the DHW computation rather than a statistical association with bleaching, it does not introduce circularity: thermal gap determines how much signal DHW *can* generate, not whether bleaching *does* occur.

### 1.5.2 4.2 Two mechanisms of DHW failure

The latitudinal gradient in this study reveals that DHW fails through two qualitatively distinct mechanisms that operate at opposite ends of the temperature spectrum.

At low latitudes (≤26°N), where median MMM was 29.81°C, summer SSTs consistently exceeded 30°C (median days30 = 9.0; 80.9% of site-years recorded ≥2 days above 30°C), yet the resulting HotSpots (SST − MMM) were small in magnitude — typically 1–2°C — because MMM itself was already near 30°C. This is a problem of metric sensitivity: the thermal stress signal exists in the environment but is compressed below the metric’s detection threshold by the elevated baseline. Corals at these sites experience chronic high-temperature exposure that drives bleaching, but DHW registers this chronic stress as a minimal departure from normal. The thermal gap — the difference between observed summer maximum SST and MMM — was only 0.928°C at low-latitude sites (median across 11 sites), setting a tight upper bound on HotSpot accumulation.

At high latitudes (>30°N), the mechanism is fundamentally different. Here, median MMM was 28.31°C, and SST rarely exceeded MMM at all: the maximum observed DHW at any high-latitude site was 2.56 °C-weeks, and no site-year exceeded even DHW ≥ 3. HotSpot events are not suppressed in magnitude — they are absent. Yet bleaching did occur at high-latitude sites (56.7% of site-years recorded >0% bleaching), likely driven by corals adapted to cooler thermal regimes for which temperatures well below 30°C — but above local tolerance limits — are sufficient to trigger stress. This is a problem of metric validity: MMM does not function as a reliable proxy for the physiological threshold above which bleaching occurs, because the temperature at which these corals experience stress may bear no fixed relationship to the climatological maximum. DHW cannot detect this bleaching in principle, because the SST values that cause stress at these sites do not exceed MMM.

Several non-exclusive hypotheses may explain why bleaching occurs at high-latitude sites without SST exceeding MMM. First, corals at these marginal sites are adapted to cooler thermal regimes with narrower thermal safety margins, meaning that even moderate warming below MMM may approach or exceed local tolerance limits. Second, high-latitude sites experience greater seasonal temperature variability, which may precondition corals to stress through mechanisms distinct from the chronic heat exposure characteristic of tropical bleaching (the thermal variability hypothesis; Safaie et al. 2018). Third, community composition at high latitudes differs markedly from low-latitude assemblages, potentially including species or genotypes with lower absolute thermal tolerance. Resolving these hypotheses requires species-level bleaching data and in-situ temperature records that are beyond the scope of the present study.

These two mechanisms — HotSpot suppression at low latitudes and sub-threshold bleaching at high latitudes — represent qualitatively different failures of the anomaly-based architecture, and they require different corrective strategies. At low latitudes, the metric’s sensitivity could be partially recovered by adjusting the HotSpot threshold below MMM or by optimising the accumulation window — the approach pursued by Whitaker and DeCarlo (2024). At high latitudes, however, no adjustment to DHW parameters can resolve the failure, because the anomaly-based framework itself is the problem: an absolute-threshold or site-specific threshold approach is required, one that does not depend on SST exceeding MMM to register stress. In the mid-latitude band, where MMM (median = 29.09°C) falls in a range that permits SST to exceed MMM by a meaningful margin during warm summers, DHW performed moderately well (AUC = 0.856 at ≥50% bleaching). The mid-latitude band represents a narrow window where the anomaly-based architecture happens to align with the thermal stress regime — not because of any design virtue, but because MMM falls in a fortuitous intermediate range.

The non-significant GEE predictor × latitude interactions reported in Section 3.2 are consistent with a signal-range interpretation: the latitudinal gradient in predictive performance is attributable to variation in the metric’s dynamic range rather than to latitudinal differences in coral thermal sensitivity. However, this interpretation should be advanced with caution. With n = 105 site-years across 21 sites, the analysis may lack sufficient power to detect moderate interaction effects, and we cannot exclude a contribution of differential coral sensitivity to the observed gradient. Future studies with larger sample sizes or species-level data would be needed to definitively partition the relative contributions of metric structure and coral biology to the observed latitudinal gradient in AUC.

### 1.5.3 4.3 Response to Whitaker and DeCarlo (2024)

The structural constraint identified in Section 4.1 provides a mechanistic explanation for the finding of Whitaker and DeCarlo (2024) that regionally optimising DHW parameters improves bleaching prediction. Their contribution was to show that a one-size-fits-all DHW definition is suboptimal. The present study explains why: regional optimisation is necessary because the information content of DHW varies systematically with MMM. In regions where MMM is high, the standard HotSpot threshold (MMM+0°C or MMM+1°C) admits only weak signals, and the optimal threshold must be lowered to recover any predictive information. In regions where MMM is low, the standard threshold may admit noise that optimisation removes. The need for regional tuning is not an empirical accident — it is a predictable consequence of anchoring a metric to a baseline that varies with climate.

This framing reinterprets regional optimisation as a symptomatic response to a structural limitation. Adjusting DHW parameters region by region can recover some predictive power, but it does not resolve the underlying problem: in warm waters, the anomaly-based architecture has a low information ceiling regardless of how parameters are tuned. At the low-latitude sites in this study, no combination of accumulation window or threshold filter would have generated DHW ≥ 4 at most sites, because the raw HotSpot values were too small. The present findings therefore complement rather than contradict Whitaker and DeCarlo (2024): their study showed that regional optimisation helps; ours shows why it is needed and where it reaches its structural limits.

### 1.5.4 4.4 Lachs versus NOAA DHW definition: an unexpected finding

A counterintuitive result emerged from the sensitivity analysis comparing two DHW formulations. The NOAA-standard DHW (which applies a MMM+1°C filter before accumulating HotSpots) yielded a higher overall AUC (0.726) than the Lachs-method DHW (MMM+0°C filter; AUC = 0.667) at the ≥50% bleaching threshold. This advantage was most pronounced at low latitudes (AUC = 0.742 vs 0.562). This result is surprising because the additional +1°C offset is more restrictive, accumulating fewer degree-days.

The likely explanation is that the +1°C filter removes small, ecologically inconsequential anomalies that act as noise in the DHW signal. At low-latitude sites where MMM ≈ 29.8°C, daily SSTs frequently exceed MMM by fractions of a degree during summer. Under the Lachs method (MMM+0°C), these minor exceedances accumulate into a DHW signal that is unrelated to bleaching — essentially, the metric detects normal summer warmth rather than anomalous stress. The NOAA +1°C filter strips out this noise, so that the remaining signal, when it occurs, reflects genuinely anomalous temperatures. The filter therefore acts as a crude signal-to-noise enhancer.

However, this improved discrimination comes at a critical operational cost. Under the NOAA definition, no site-year in the entire monitoring network reached DHW ≥ 4 — not a single alert would have been issued across five years and 113 site-years, during which 65 site-years recorded bleaching. The NOAA definition thus presents a paradox: it improves the continuous discriminatory performance of the DHW signal while simultaneously rendering the threshold-based alert system completely non-functional. This distinction between DHW as a continuous predictor and DHW as an alert system deserves emphasis: the two are separate questions, and the latter — which determines whether management actions are triggered — is the more operationally consequential.

### 1.5.5 4.5 The case for absolute temperature metrics

The consistent superiority of days30 across latitudinal bands and bleaching severity thresholds (overall ΔAUC = 0.260 at ≥50% bleaching, p < 0.001) arises from its independence from MMM. By counting days above a fixed absolute temperature, days30 measures proximity to a physiologically meaningful boundary regardless of local climatology. At low-latitude sites, where DHW was near-uninformative (AUC = 0.562), days30 achieved an AUC of 0.845 — a performance gap of 0.293 that reflects the fundamental structural difference between anomaly-based and absolute-threshold metrics.

The advantage of absolute-threshold metrics, however, should not be overstated. The 30°C threshold was identified as optimal for this dataset by Fukui (2026), and its applicability to other regions is untested. At high-latitude sites in this study, days30 performance was high (AUC = 0.964 at ≥50% bleaching), but this reflected the metric’s ability to separate the few high-latitude sites that experienced SST ≥ 30°C from those that did not — a separation that may be confounded with latitudinal position rather than indicative of 30°C as a universal physiological threshold. More importantly, at the >0% bleaching threshold in the high-latitude band, days30 AUC was only 0.618 — near chance-level performance — indicating that low-severity bleaching at high-latitude sites occurs at temperatures below 30°C and is not captured by this threshold. This limitation is not incidental: at high latitudes, the >0% bleaching threshold captures stress events that occur below both the MMM-relative anomaly range (rendering DHW insensitive) and the 30°C absolute threshold (rendering days30 insensitive). The coexistence of metric failure for both DHW and days30 in this band underscores that no single thermal metric — whether anomaly-based or absolute-threshold — can adequately predict bleaching across the full latitudinal range of Japanese coral reefs. This structural incompleteness, rather than the superiority of any one metric, is the most important practical conclusion of this study. These findings point toward the need for regionally calibrated absolute thresholds — or, more ambitiously, site-specific thresholds derived from MMM-relative reasoning — a direction we leave to subsequent work.

### 1.5.6 4.6 Limitations

Six limitations qualify these findings.

First, this study analysed 26 sites in a single country. While the latitudinal gradient provides a controlled test of DHW’s structural properties, generalisation to other subtropical and tropical reef regions — where species composition, oceanographic regimes, and bleaching histories differ — requires independent validation.

Second, all thermal metrics were computed from satellite SST (MUR v4.1), which does not capture fine-scale temperature variability at the reef scale (depth, flow-mediated cooling, diurnal stratification). Validation against in-situ temperature records is deferred to subsequent work.

Third, the five-year observational window (2020–2024) is too short to capture decadal-scale shifts in MMM under climate change. If MMM is itself shifting upward, the DHW values computed here may underestimate the long-term degree of HotSpot suppression. This bias is conservative with respect to our central argument: a rising MMM would further compress the thermal gap and exacerbate DHW’s structural insensitivity.

Fourth, species composition varies across the latitudinal gradient, and the higher bleaching prevalence at low-latitude sites could partly reflect greater representation of thermally sensitive taxa (e.g., *Acropora*). This confound was discussed in Fukui (2026) and cannot be resolved without species-level bleaching data.

Fifth, the MMM climatology was computed over 2019–2024 rather than the multi-decadal baseline (1985–1993) used by Lachs et al. (2021) and the 1985–2012 period used by NOAA Coral Reef Watch. The short, recent baseline captures the contemporary thermal regime including recent warming. This means that our MMM values are likely higher than those derived from historical baselines, which would make our DHW values more conservative (lower) than those computed from historical climatologies. This bias again works against our central finding — with a lower (historical) MMM, DHW would have somewhat greater dynamic range, and its performance would be somewhat better than reported here. A supplementary comparison with the NOAA CRW operational product (Fig. S2) confirmed this prediction empirically: CRW DHW, anchored to the 1985–2012 climatology, reached the ≥4 alert threshold in 50.5% of site-years, compared with only 3.6% under our contemporary-baseline calculation. Although the two products differ in spatial resolution (MUR 0.01° vs CRW 0.05°), accumulation window (8 vs 12 weeks), and HotSpot threshold (MMM+0°C vs MMM+1°C), the dominant driver of the discrepancy is the baseline period: CRW’s historical MMM is lower, producing larger HotSpots and higher DHW values. This comparison demonstrates that our MUR-derived results are conservative relative to the operational product, and that the structural insensitivity documented here reflects the progressive erosion of DHW signal range as MMM rises with climate change. Construction of an independent multi-decadal baseline from MUR SST is constrained by the dataset’s start date (2002), and a sensitivity analysis recomputing MMM over 2002–2024 is a priority for subsequent work.

Sixth, the GEE predictor × latitude interactions were non-significant (DHW: p = 0.183; days30: p = 0.607), but the balanced panel (n = 105 across 21 sites) may lack statistical power to detect moderate interaction effects. The ΔAUC gradient across latitudinal bands was clear and consistent in direction (largest at low latitudes, smallest at high latitudes), even though the formal interaction test did not reach significance.

### 1.5.7 4.7 Implications for monitoring and management

The standard NOAA Coral Reef Watch alert thresholds (DHW ≥ 4 and DHW ≥ 8) are structurally non-functional across Japan’s coral monitoring network. Across 113 site-years encompassing two major bleaching events, DHW ≥ 4 detected only 6.2% of bleaching occurrences. Under the NOAA-standard DHW definition (MMM+1°C filter), no site-year reached DHW ≥ 4 at all. These are not marginal failures of a threshold that needs minor adjustment; they reflect a structural mismatch between the metric’s design and the thermal environment.

For regional monitoring programs operating in warm subtropical waters — where MMM approaches or exceeds 29°C — these findings suggest that the anomaly-based DHW framework cannot serve as a reliable early warning system regardless of how alert thresholds are calibrated. Alternative approaches that do not depend on MMM-relative anomalies — whether absolute-temperature metrics, composite indices incorporating non-thermal variables, or machine-learning approaches that learn region-specific nonlinearities — deserve systematic evaluation. The development of margin-aware metrics that explicitly account for the distance between climatological baselines and physiological tolerance limits is a promising direction that we pursue in subsequent work.

More broadly, the structural insensitivity documented here may apply to any reef region where recent warming has pushed MMM close to coral thermal tolerance limits. As ocean temperatures continue to rise, the thermal gap available for HotSpot accumulation will narrow further across an expanding proportion of global reef areas. The anomaly-based architecture thus contains a built-in convergence toward insensitivity under sustained warming — a structural property that current regional optimisation efforts cannot overcome. The global domain in which DHW retains adequate signal range will progressively narrow — a consequence of the anomaly-based architecture’s dependence on a baseline that tracks the very warming trend it is meant to detect. Japan’s latitudinal gradient, spanning from equatorial-like thermal regimes to marginal temperate reefs under a single monitoring protocol, offers an early window into this global trajectory. Notably, the spatial pattern documented here converges with the temporal modelling of Lachs et al. (2023), who showed that simulated increases in coral thermal tolerance at Palau — which effectively raise the local MMM — progressively eliminated the DHW signal over decades. Our cross-sectional analysis demonstrates that this same signal erosion is already manifest in the spatial domain: sites with higher contemporary MMM exhibit the same DHW insensitivity that Lachs et al. projected for the future under continued warming. Together, these independent lines of evidence — one temporal, one spatial — indicate that DHW’s structural insensitivity is not a local calibration problem but a general consequence of any process that narrows the gap between the climatological baseline and the thermal stress threshold. The Monitoring Site 1000 dataset demonstrates that standardised, long-term monitoring networks can reveal structural limitations in global metrics that aggregate analyses across heterogeneous protocols cannot detect.

## 1.6 Acknowledgements

The author thanks the Ministry of the Environment, Japan, and the Biodiversity Center of Japan for access to the Monitoring Site 1000 coral reef survey data (fiscal years 2020–2024), approved under permit Kansei-Tahatsu No. 2602271. The author also thanks K. Kozono at the Biodiversity Center for administrative support.

## 1.7 Data Availability

The Monitoring Site 1000 coral reef survey data are managed by the Biodiversity Center of Japan, Ministry of the Environment. Access requests should be directed to the Biodiversity Center. MUR SST data are publicly available from NASA JPL (https://podaac.jpl.nasa.gov/). Analysis code is available at https://github.com/hi-rokifukui/marine-research.

## 1.8 Author Contributions

HF conceived the study, conducted all analyses, and wrote the manuscript.

## 1.11 Supplementary Material

**Fig. S1:**
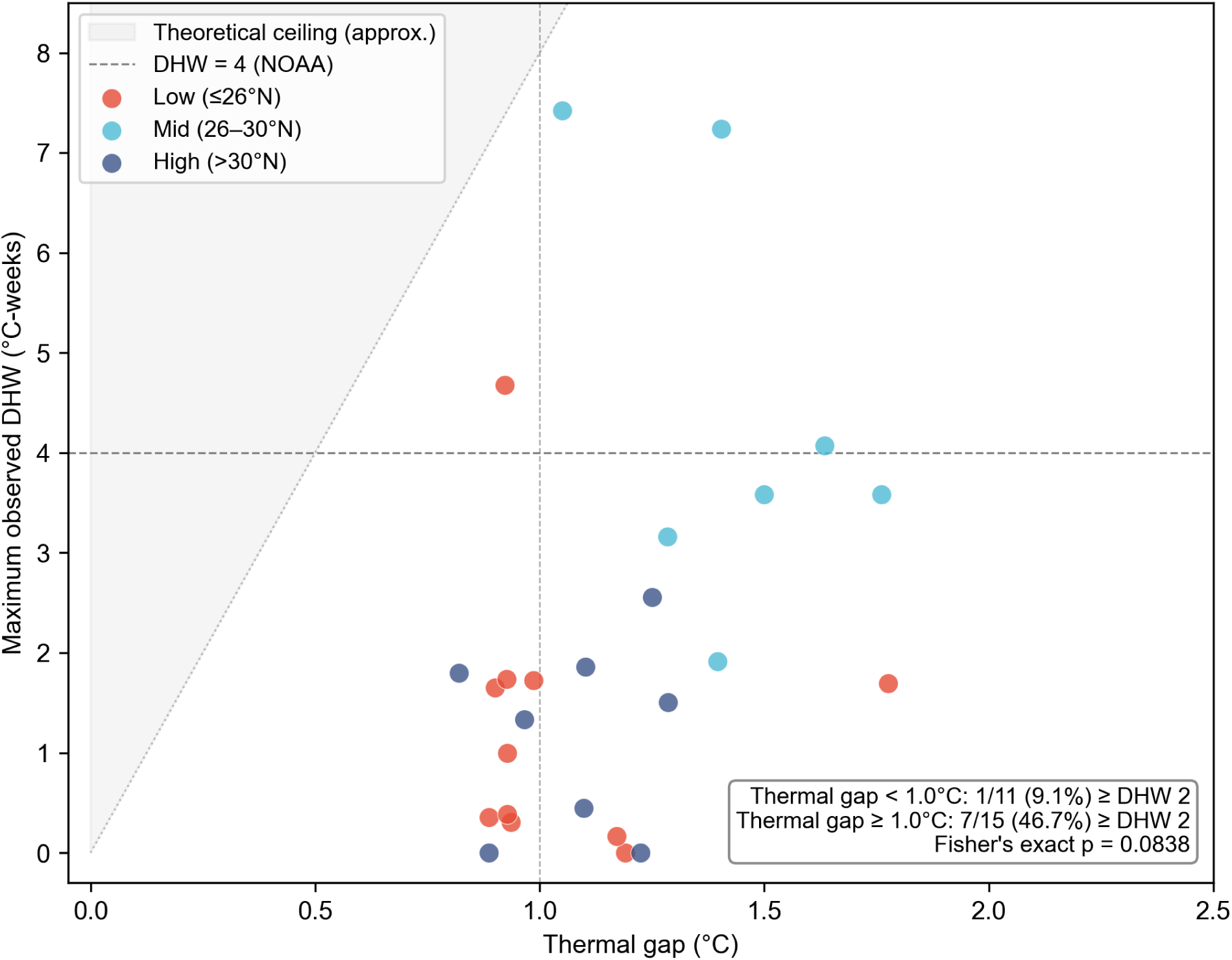
Relationship between thermal gap (summer maximum SST − MMM) and maximum observed DHW across 26 sites. Points are coloured by latitudinal band. Horizontal dashed line indicates the DHW ≥ 4 alert threshold. Vertical dashed line at 1.0°C marks the boundary used for the Fisher’s exact test (see Section 3.3). The shaded region represents the approximate theoretical ceiling under constant exceedance (thermal gap × 8 weeks).

**Fig. S2:**
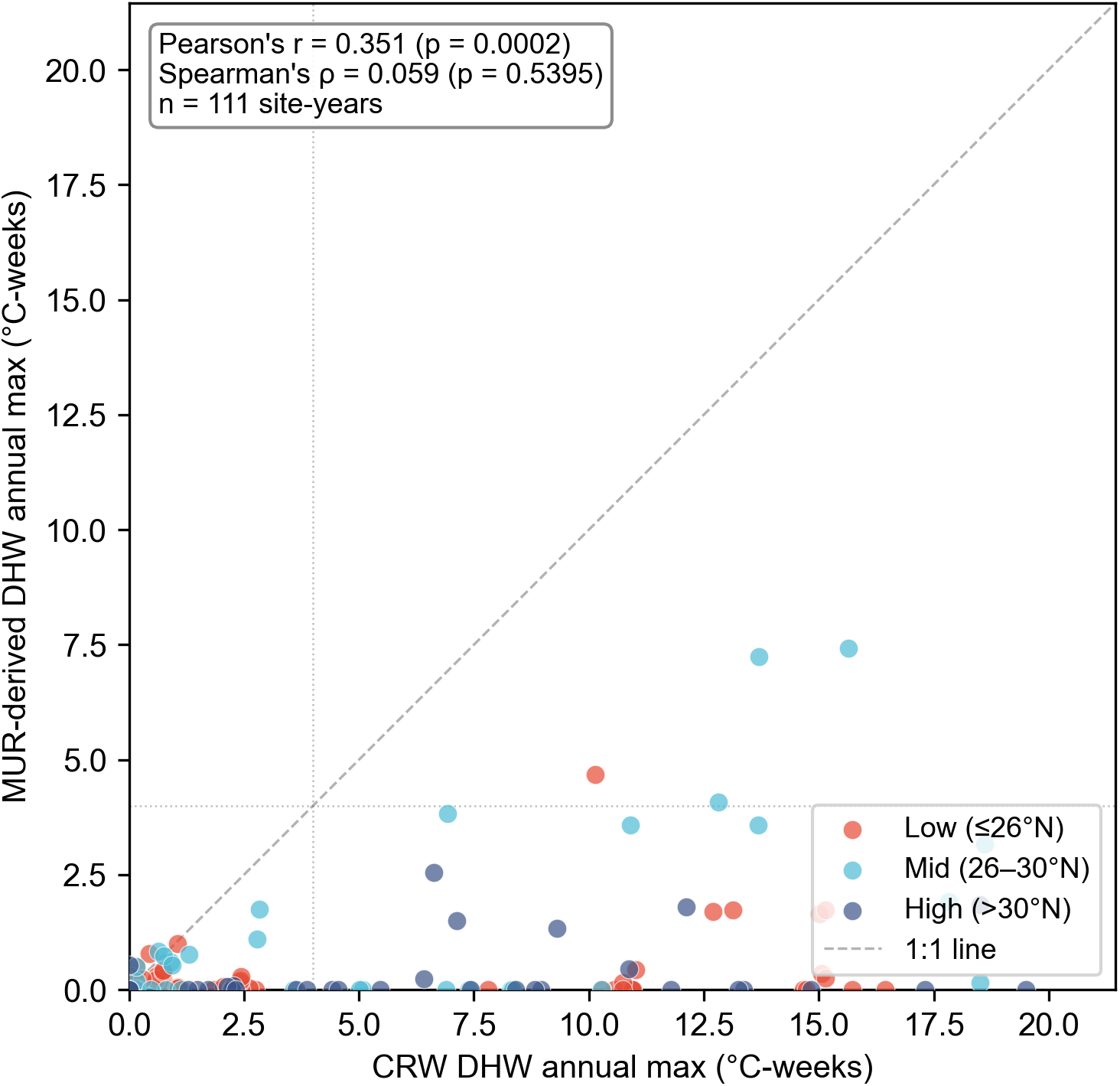
Comparison of annual maximum DHW between MUR-derived values (this study; Lachs method, 8-week window, MMM from 2019–2024) and NOAA CRW operational product (v3.1; 12-week window, MMM+1°C threshold, 1985–2012 climatology). Each point represents one site-year (n = 111). Dashed diagonal line indicates the 1:1 relationship. Dotted lines at DHW = 4 mark the standard alert threshold on each axis. Points are coloured by latitudinal band. CRW DHW values are systematically higher than MUR-derived values, reflecting the lower historical MMM baseline.

