## Supplementary figures and images for "1 Degree heating weeks fail to reach alert thresholds yet coral bleaching is widespread: structural insensitivity of anomaly-based metrics across Japan’s latitudinal gradient"

### Figure S1

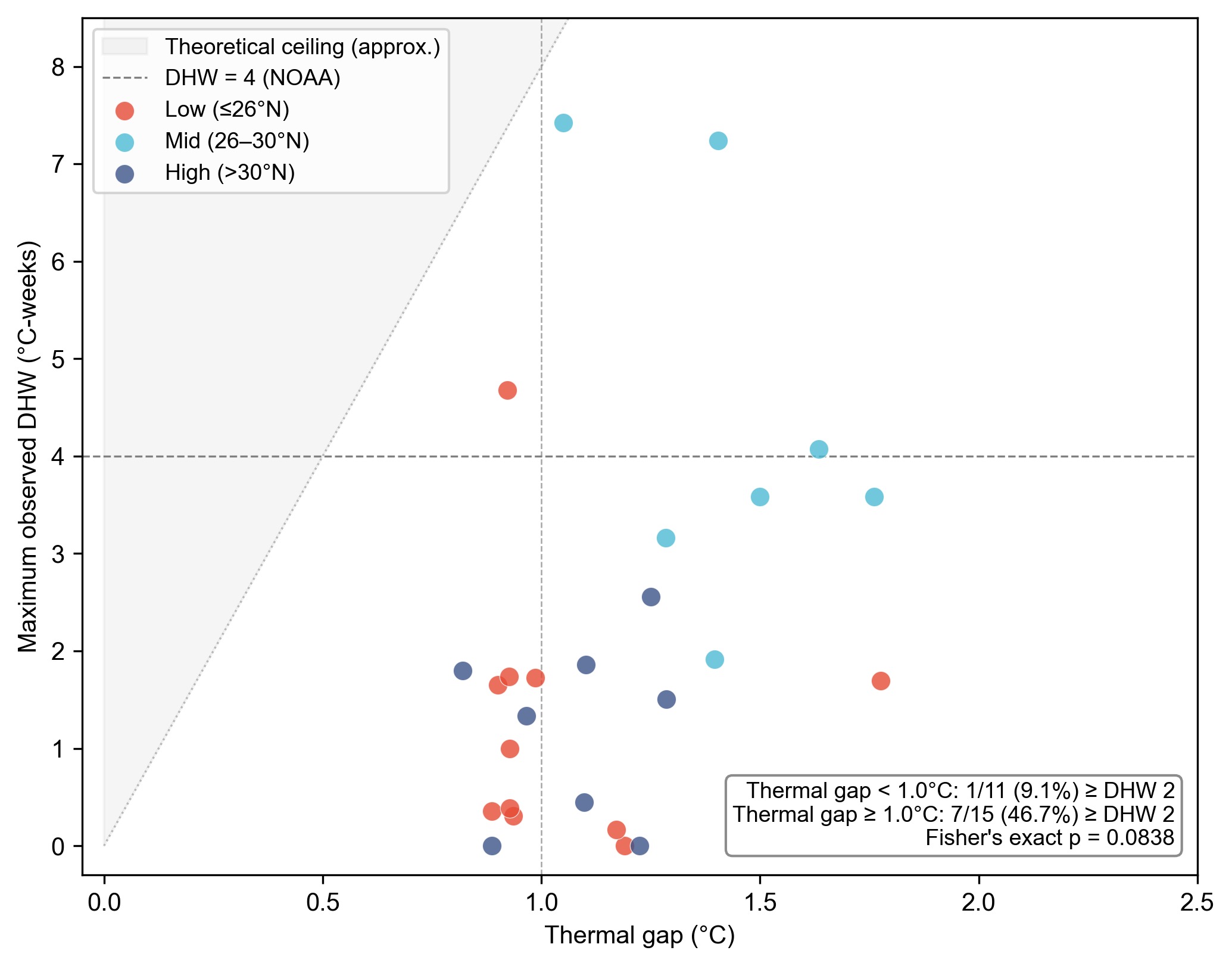

### Figure S2

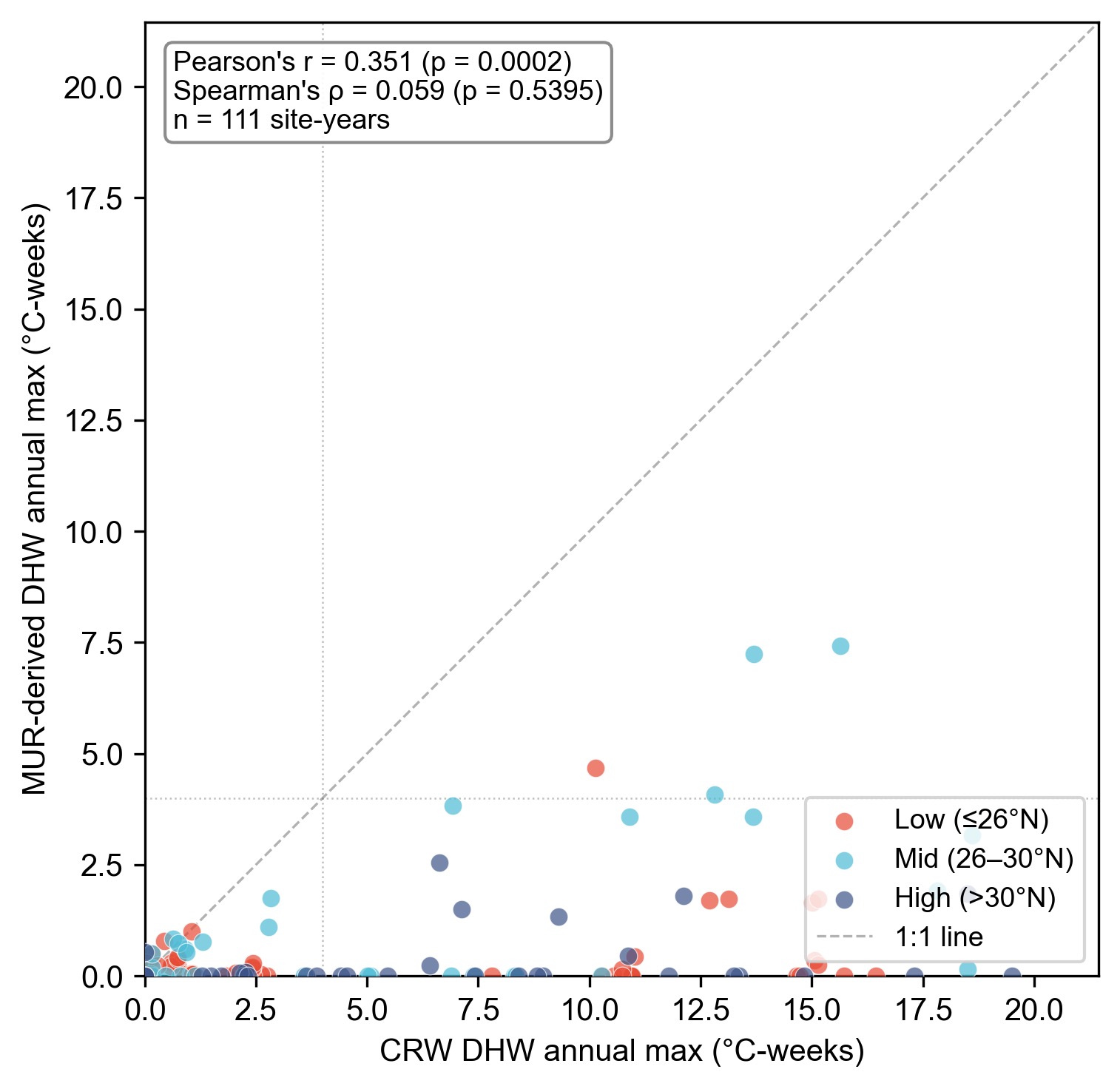
